# Electroacupuncture promotes post-stroke motor recovery through miR-124-3p-mediated regulation of NRG1/ErbB4 signaling pathway

**DOI:** 10.1101/2025.05.27.656264

**Authors:** Debiao Yu, Peng Chen, Xiaoting Chen, Yaoyu Lin, Fa Lin, Nan Chen, Fuchun Wu, Bin Shao

## Abstract

**Objective:** Electroacupuncture has demonstrated beneficial effects in post-stroke motor dysfunction, yet the molecular mechanisms underlying its therapeutic efficacy remain incompletely understood. This study aims to investigate whether electroacupuncture promotes post-stroke motor function recovery by modulating the interaction between miR-124-3p and the NRG1/ErbB4 signaling pathway, specifically exploring whether miR-124-3p directly targets NRG1 to regulate neural plasticity in a focal cerebral ischemia rat model.

**Methods:** The ischemic stroke model was established by middle cerebral artery occlusion/reperfusion (MCAO/R) in adult rats. These rats were randomly divided into sham, model, electroacupuncture and model plus miR-124-3p inhibitor groups. The model group, electroacupuncture group and model plus miR-124-3p inhibitor group received EA intervention 24 h after modelling for 7 consecutive days. Behavioural function was assessed by Zea Longa score and mechanical pain rating. Hippocampal damage was detected by HE staining and neuronal apoptosis was observed by TUNEL staining. IL-1β and IL-18 levels were measured by ELISA. PCR and Western blotting were used to detect the expression of miR-124-3p and inflammatory pathway proteins. The interaction between miR-124-3p and TLR4 was verified by dual-luciferase reporter assay.

**Results:** Electroacupuncture improved motor function in rat model of MCAO/R, as evidenced by improved Zea Longa scores and decreased mechanical withdrawal thresholds. Electroacupuncture significantly attenuated neuronal damage and also inhibited the inflammatory response by decreasing IL-18 and IL-1β levels (P < 0.001). Notably, electroacupuncture upregulated miR-124-3p expression (P < 0.0001) and activated the NRG1/ErbB4 signaling pathway in the hippocampus. When miR-124-3p was inhibited, NRG1 protein expression decreased while GABA expression tended to increase. Dual-luciferase reporter assays confirmed that miR-124-3p directly targets the 3’UTR of NRG1 mRNA and regulates its expression at the translational level. These findings suggest that electroacupuncture may alleviate neuronal and axonal damage by modulating miR-124-3p/NRG1/ErbB4 signalling and regulating GABA release.

**Conclusion:** Electroacupuncture can ameliorate motor dysfunction induced after brain I/R injury by targeting and modulating the NRG1-ErbB4 signaling pathway via miR-124-3p. These data are expected to provide new insights into the mechanisms of electroacupuncture for the prevention of potential targets for the recovery of motor dysfunction after stroke.

## 1 Introduction

Stroke is the third leading cause of death and disability worldwide^1,2^, with over 80% of patients experiencing impaired limb motor function, significantly impacting their quality of life^3^. Current intervention approaches such as conventional rehabilitation therapy, constraint-induced movement therapy, and pharmacological treatments often face challenges including limited efficacy, prolonged treatment duration, and poor patient adherence, creating an urgent need to explore more effective therapeutic strategies^4,5^.

Recovery of motor function after stroke is primarily based on neuroplasticity mechanisms^6,7^. Neuroplasticity refers to the brain’s ability to reorganise its structure and function, making new neural connections to compensate for damaged functions. This plasticity provides the critical structural basis for motor recovery after stroke^8^. At the molecular level, synaptic plasticity involves the precise regulation of multiple signaling pathways^9,10^. Recent studies have shown that microRNAs (miRNAs) play a central role in this process^11^. miR-124-3p, as one of the most abundantly expressed miRNAs in the central nervous system, has significant effects on synaptic plasticity^12^. Several studies have confirmed that post-stroke changes in miR-124-3p expression levels are closely associated with recovery of neurological function, primarily through the targeted regulation of various genes involved in synaptic plasticity^13,14^.

Meanwhile, the signaling pathway formed by neuregulin-1 (NRG1) and its receptor ErbB4 plays a crucial role in the development and maintenance of the neural system by regulating neuronal migration, synapse formation, myelination and synaptic transmission^15^. During recovery from stroke, activation of the NRG1-ErbB4 pathway promotes axonal regeneration, synaptic plasticity and restoration of neural function^16^. Notably, bioinformatic analysis has identified NRG1 as a potential direct target of miR-124-3p, a finding validated in models of myocardial ischemia^17^. Given the commonalities in ischemic response mechanisms between myocardial and neural tissues, this provides a theoretical basis for exploring the role of the miR-124-3p-NRG1/ErbB4 pathway in cerebral ischemia.

Acupuncture, a characteristic therapy of Traditional Chinese Medicine, has been recognised by the World Health Organisation as an effective complementary intervention for the treatment of stroke^18^. Electroacupuncture, a modernised technological derivative of acupuncture, provides a reliable platform for basic research and clinical applications through precise and controllable stimulation^19^. Recent studies have shown that electroacupuncture can reduce neuronal apoptosis by regulating miR-124-3p expression, inhibiting GSK-3β activity and subsequently reducing Cyp-D-mediated mitochondrial permeability transition pore opening^20^. At the same time, research suggests that electroacupuncture can activate the NRG1-ErbB4 signaling pathway, promoting neuroplasticity and significantly improving neurological function after cerebral ischemia^15^. These findings suggest that electroacupuncture may simultaneously act on both the miR-124-3p and NRG1-ErbB4 signaling pathways. However, whether there is a cross-regulatory relationship between these two pathways, in particular the specific molecular mechanisms by which electroacupuncture exerts therapeutic effects by coordinating the interaction between miRNA and protein pathways, remains to be elucidated.

Based on the above research background, we hypothesise that: (1) miR-124-3p may directly target NRG1 to regulate the NRG1/ErbB4 signaling pathway, thereby influencing neuroplasticity and functional recovery; (2) electroacupuncture may indirectly modulate the NRG1/ErbB4 signaling pathway by regulating miR-124-3p expression, thereby promoting motor function recovery after stroke. This study will establish a focal cerebral ischemia model in rats to systematically investigate this mechanism, which will not only contribute to a deeper understanding of the molecular basis of neuroplasticity after stroke, but also provide new theoretical evidence for electroacupuncture treatment of stroke and potentially provide insights for the development of novel intervention strategies based on miR-124-3p.

## 2 Materials and Methods

### 2.1 Animals

Twenty-four Sprague-Dawley (SD) rats (weighing 250-300 g) were purchased from SPF (Beijing) Biotechnology Co., Ltd., license number: SCXK (Beijing) 2019-0010. The rats were housed under controlled conditions of temperature (20-26°C), humidity (40-70%), and a 12h/12h light/dark cycle. All rats were acclimatized for one week prior to the experiment. This study was approved by the Animal Ethics Committee of Fujian Provincial Hospital Affiliated to Fuzhou University (approval number: LL-202306130001). All animal procedures were conducted in accordance with the ARRIVE guidelines and the International Guidelines for the Care and Use of Laboratory Animals.

### 2.2 Preparation of MCAO/R rat models and group

After anaesthesia with intraperitoneal injection of 10% chloral hydrate (350 mg/kg), rats were fixed on a stereotaxic apparatus (RWD Life Science, China) for stereotaxic injection (left brain, -0.8 mm posterior, -1.5 mm lateral, -4.5 mm deep). According to the above grouping, rats requiring drug treatment received miR124-3p (UAAGGCACGCGGUGAAUGCC) antagonist at a dose of 10 nmol per rat, while the other groups received an equal volume of saline. The middle cerebral artery occlusion/reperfusion (MCAO/R) model was established one day after injection. In the MCAO/R model, the external carotid artery (ECA) and its branches were first ligated, then a nylon filament was inserted through the ECA into the internal carotid artery to reach the anterior cerebral artery (ACA), mechanically occluding the blood supply to the middle cerebral artery (MCA) for 120 minutes. The sham surgery group underwent the same procedure but without the nylon thread. The Zea Longa scoring method was used to verify the success of the model, with scores of 1-4 indicating an effective MCAO/R model.

After a one-week acclimatization period, the experimental animals were randomly divided into four groups (n=10 per group) using Random Allocation Software 2.0: sham group (control), model group, electroacupuncture group, and model plus miR-124-3p inhibitor group. A double-blind design was implemented throughout the experiment, with grouping and interventions performed by investigators who were not involved in subsequent experimental evaluations. All scoring, histological analyses, and biochemical tests were performed by investigators blinded to group allocation.

### 2.3 Electroacupuncture intervention

Electroacupuncture was administered the day after modelling. Acupoints Zusanli (ST36) and Quchi (LI11) were selected for electroacupuncture treatment of ischemic stroke in the affected limb. The rat acupoints were located according to the “Standard for Experimental Animal Acupoints”. LI11 is located in the depression on the medial side of the extensor carpi radialis, at the lateral end of the elbow crease; ST36 is located on the craniolateral surface of the lower leg at approximately one fifth of the distance from the proximal end, at the distal end of the tibia, in the depression between the cranial tibial muscle and the long digital extensor muscle. Acupuncture needles (0.16×7 mm) were inserted to a depth of 2 mm and connected to a G6805-1I electro-acupuncture therapeutic device. The stimulation intensity was set to 1 mA at a frequency of 2 Hz. Electroacupuncture treatment was given once a day for 20 minutes for 5 consecutive days. Zea longa scoring was performed twice to assess neurological deficit: once after modelling surgery (before electroacupuncture) and once after electroacupuncture. All mice in the other groups were subjected to the same handling and restraint procedures.

### 2.4 Neurological Deficit Scoring

1. Zea Longa neurological score: Neurological deficits were scored according to the following criteria: 0 points: no neurological deficits; 1 point: inability to fully extend the right forelimb; 2 points: circling to the right during free movement; 3 points: falling to the right during free movement; 4 points: inability to walk spontaneously or loss of consciousness.
2. Mechanical pain rating: Mechanical pain thresholds were quantified using von Frey filaments 24 hours after the end of treatment. Paw withdrawal thresholds were measured to quantify the pain response in the rats.

### 2.5 Tissue Collection and Sample Processing

After surgery, rats were anaesthetised with 10% chloral hydrate (40 mg/kg) administered by intraperitoneal injection, the dose being calculated on the basis of body weight. Subsequent tissue collection and sample processing were performed according to different experimental requirements. Rats were anaesthetised and hippocampal tissue was rapidly isolated. The whole hippocampus was fixed in 4% paraformaldehyde for 24 hours. The samples were then rinsed with PBS, dehydrated through an ethanol gradient, cleared with xylene and finally embedded in paraffin for subsequent HE staining and TUNEL detection. For ELISA and Western blot analyses, ELISA samples were snap frozen in liquid nitrogen and then stored at -80°C. Western blot samples were immediately homogenised in pre-cooled tissue lysis buffer, centrifuged at 4°C and the supernatant collected, aliquoted and stored at -80°C. For dual luciferase detection, cell lysates were extracted from cultured chondrocytes and immediately subjected to luciferase activity measurement.

### 2.6 Hematoxylin–eosin (HE) and Nissl Staining

The hippocampus of each rat was fixed with 4% paraformaldehyde (Beyotime Biotechnology Co., Shanghai, China) for 48 h at 4 °C. After dehydrating the hippocampal samples and embedding them in paraffin, 5-µm thick slices were prepared. Then, the slices were treated with xylene and gradient alcohol to dewax and rehydrate them, respectively.

For HE staining, hematoxylin and eosin solutions (both from Lianmai Biological Engineering Co., Shanghai, China) were sequentially added to the sections for staining for 5 and 2 min, respectively. For Nissl staining, the sections were stained with Nissl solution (Yiyan Biotechnology Co., Shanghai, China) for 5 min. After dehydration using gradient alcohol and transparency using xylene, the dried sections were sealed with neutral resin for histopathological and neuronal observations under an optical microscope (CX41, Olympus, Tokyo, Japan).

### 2.7 Enzyme-linked immunosorbent assay(ELISA)

ELISA was used to determine the levels of IL-1β and IL-18. Rat serum was collected and centrifuged at 1000g for 20 min. The assay was performed according to the manufacturer’s instructions (ExCell Biology, Shanghai, China). The specific procedure was as follows The ELISA kit was equilibrated at room temperature. Standards were prepared according to the manufacturer’s instructions. Blank, standard and sample wells were prepared. No samples or enzyme-labelled reagents were added to the blank wells. 50 μL of sample or standard was added to the remaining wells, followed by 100 μL of enzyme-labelled reagent. The plate was incubated for 1 hour at 37°C. After five washes, chromogenic substrate was added and incubated at 37°C for 15 minutes. The reaction was stopped by adding stop solution and the absorbance of each well was measured at a wavelength of 450 nm.

### 2.8 Real-Time Quantitative Reverse Transcription Polymerase Chain Reaction (qRT-PCR)

Total RNA was extracted from rat brain tissue using Trizol reagent (Invitrogen, USA), while miRNA was isolated using a miRNA purification kit (CW0627S, CWBIO, China). RNA concentration and purity were measured using an NP80 UV spectrophotometer (NanoPhotometer, Germany), and samples with OD260/OD280 ratios between 1.8-2.0 were considered acceptable. miRNA was reverse transcribed into cDNA using miRNA 1st Strand cDNA Synthesis Kit (by stem-loop) (MR101-02, Vazyme, China). qPCR reactions were performed using miRNA Universal SYBR qPCR Master Mix (MQ101-02, Vazyme, China) on a CFX Connect™ fluorescence PCR system (Bio-Rad, USA). The reaction protocol was as follows: pre-denaturation at 95°C for 10 min; denaturation at 95°C for 10 s; annealing at 58°C for 30 s; extension at 72°C for 30 s; for 40 cycles. Each sample was analyzed in triplicate using U6 as an internal reference. Relative gene expression was calculated using the 2^-ΔΔCT method. Primer sequences are listed in the Table1 below.

### 2.9 Western blot (WB)

Rat hippocampal tissue was collected and homogenised in RIPA lysis buffer containing protease and phosphatase inhibitors using a tissue homogeniser to extract total protein. The homogenate was centrifuged at 12,000 rpm for 10 min at 4°C and the supernatant was collected. Protein quantification was performed using a BCA protein assay kit. After protein denaturation, samples were separated by sodium dodecyl sulfate-polyacrylamide gel electrophoresis (SDS-PAGE) for 1.5 h. Proteins were transferred to PVDF membranes (Millipore) at a constant current of 300 mA for different durations: 1 h for NRG1, 2 h for ErbB4, TLR4 and NLRP3 and 0.5 h for ASC. The PVDF membranes were blocked with nonfat milk and then incubated with primary antibodies overnight at 4°C. The next day, the membranes were incubated with secondary antibodies for 2 h at room temperature. Membranes were wetted with chemiluminescent reagent and visualised using a high sensitivity chemiluminescence imaging system. Band intensities were analysed using Image J software, with β-actin used as an internal reference to calculate the relative expression of target proteins.

### 2.10 Analysis of the dual-luciferase reporter gene

Wild-type and mutant TLR4 3’UTR reporter plasmids were constructed using pmirGLO vector backbone (Promega, USA). 293T cells (BNCC353535, BNBIO, China) were seeded in 24-well plates and transfected at 70-80% cell confluence using Lipofectamine 3000 (Invitrogen, USA) according to the manufacturer’s instructions. Each well was transfected with a combination of 200 ng wild-type or mutant TLR4-3’UTR reporter plasmid and 50 nM miR-124-3p agomir or negative control (RiboBio, China). Cells were harvested 48 h post-transfection and 200 μL lysis buffer was added to each well. After 20 min lysis at 4°C, 70 μL of cell lysate was transferred to a black 96-well plate. A dual-luciferase reporter assay kit (RG027, Beyotime, China) was used according to the manufacturer’s protocol: firefly luciferase substrate (100 μL/well) was added first to measure firefly luciferase activity, followed by renilla luciferase substrate (100 μL/well) to measure renilla luciferase activity. Luminescence was detected using a SuPerMax 3100 automated microplate reader (Flash Spectrum, China), and firefly luciferase activity was normalised to renilla luciferase activity.

### 2.11 Statistical analysis

Statistical analysis was performed using IBM SPSS Statistics version 26.0 (IBM Corp, USA). Data are presented as mean ± standard deviation (±s). Normality of data distribution was assessed using the Shapiro-Wilk test, while homogeneity of variance was assessed using Levene’s test. For data with normal distribution and homogeneous variance, one-way analysis of variance (ANOVA) was used for multiple group comparisons, followed by the least significant difference (LSD) test for pairwise comparisons. Non-normally distributed data were analyzed using the non-parametric Kruskal-Wallis test, with the Mann-Whitney U test for between-group comparisons. All statistical analyses were performed using two-tailed tests, with *P*<0.05 considered statistically significant. GraphPad Prism 10.0 software (GraphPad Software, USA) was used to generate the figures.

## 3 Results

### 3.1 Electroacupuncture restores motor function in MCAO/R rats

The effects of electroacupuncture treatment on ischemic stroke-induced motor dysfunction were evaluated using the Zea Longa neurological deficit score and the mechanical withdrawal threshold test. As shown in Figure 1A and Figure 1B, rats in the model group showed decreased motor function and increased pain threshold compared with the blank group, indicating successful early modeling. After 5 days of electroacupuncture treatment in MCAO/R rats, improved motor function and reduced pain threshold were observed in the model plus electroacupuncture group compared with the model group.

**Fig 1.**
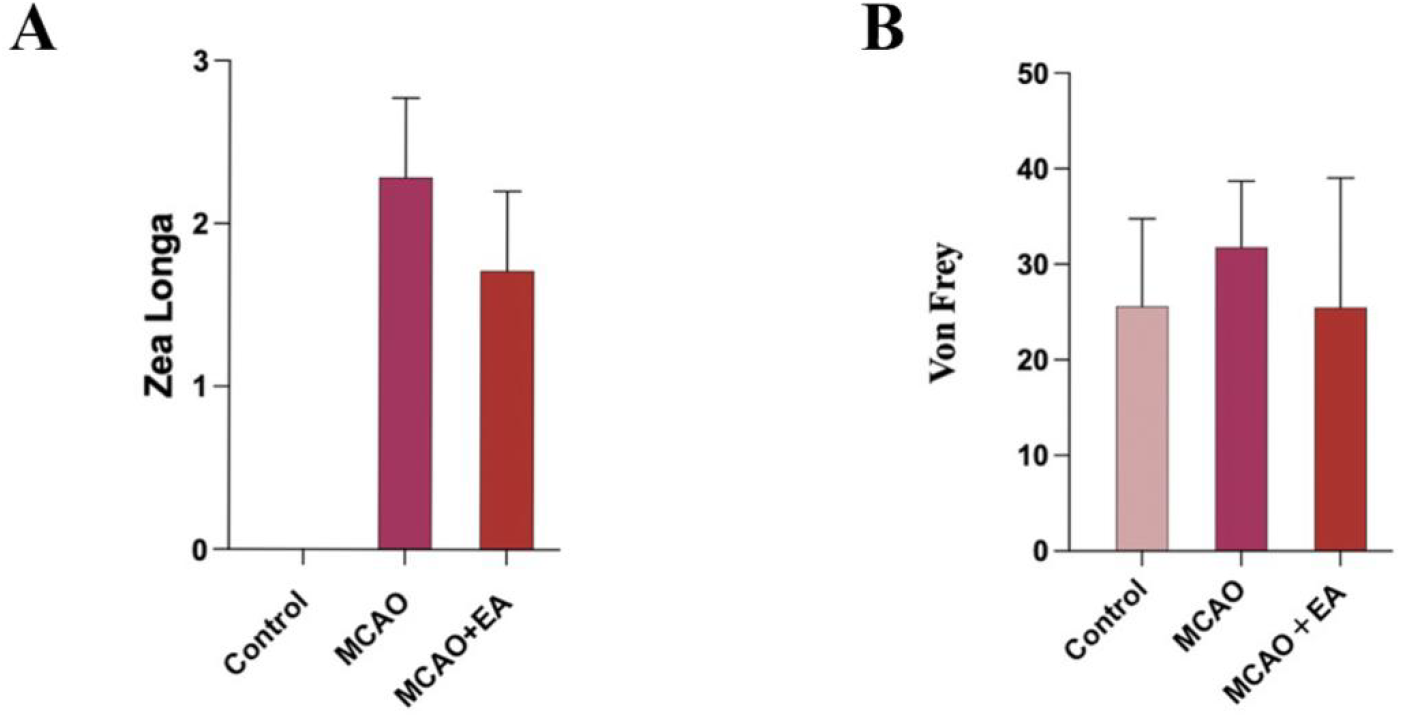

### 3.2 Electroacupuncture ameliorates neuronal damage and inflammatory response induced by MCAO/R

The results of H&E and Nissl staining (Figure 2A) showed that the blank group had clear and intact structures with neurons arranged in an orderly fashion. In the model group, neuronal cells showed nuclear pyknosis, vacuolation and even cell death, with abnormally shaped deep blue Nissl bodies visible in Nissl staining. Compared with the model group, the model plus electroacupuncture group showed significantly improved neuronal cell damage and reduced abnormally shaped deep blue Nissl bodies. These results suggest that electroacupuncture treatment can alleviate MCAO/R-induced neuronal damage. To determine the effect of electroacupuncture on MCAO/R-induced inflammatory response, we examined inflammatory factors including IL-18 and IL-1β. The results showed that IL-18 and IL-1β levels were elevated after ischemia-reperfusion (*P* < 0.0001), while electroacupuncture treatment inhibited the elevation of IL-18 and IL-1β (*P* < 0.0001, *P* < 0.001, respectively) (Figure 2B, Figure 2C).

**Fig 2.**
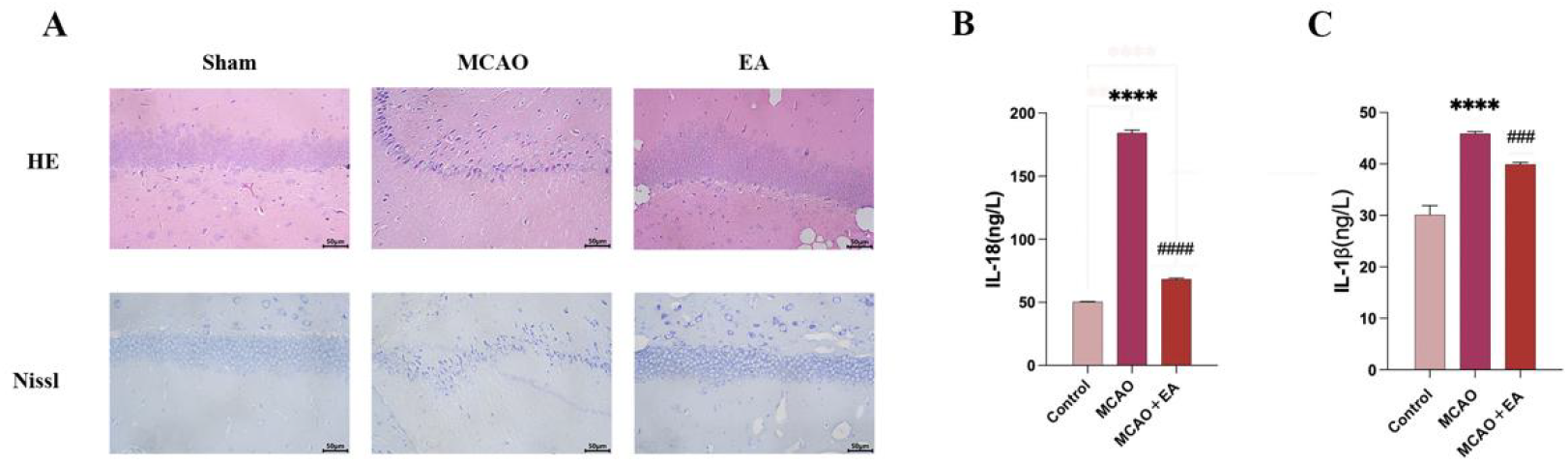

### 3.3 Electroacupuncture regulates miR-124-3p levels and activates the NRG1/ErbB4 signaling pathway in the hippocampus of MCAO/R rats

Since miR-124-3p serves as a potential biomarker of ischemic brain injury and plays a role in promoting recovery of neurological function, we investigated its expression. Results showed that miR-124-3p was significantly upregulated in the model plus electroacupuncture group (*P* < 0.0001) (Figure 3A). To further investigate whether the downstream pathways of miR-124-3p were also regulated by electroacupuncture, we examined the protein expression of NRG1 and ErbB4 in the hippocampus. The results showed that compared with the control group, the protein levels of NRG1 and ErbB4 were significantly decreased in the model group (*P* < 0.05), while electroacupuncture upregulated the protein levels of NRG1 and ErbB4 (P < 0.05). In the model plus inhibitor group, NRG1 levels decreased (*P* < 0.05), but ErbB4 levels increased (*P* < 0.05) (Figure 3B, Figure 3C).

**Fig 3.**
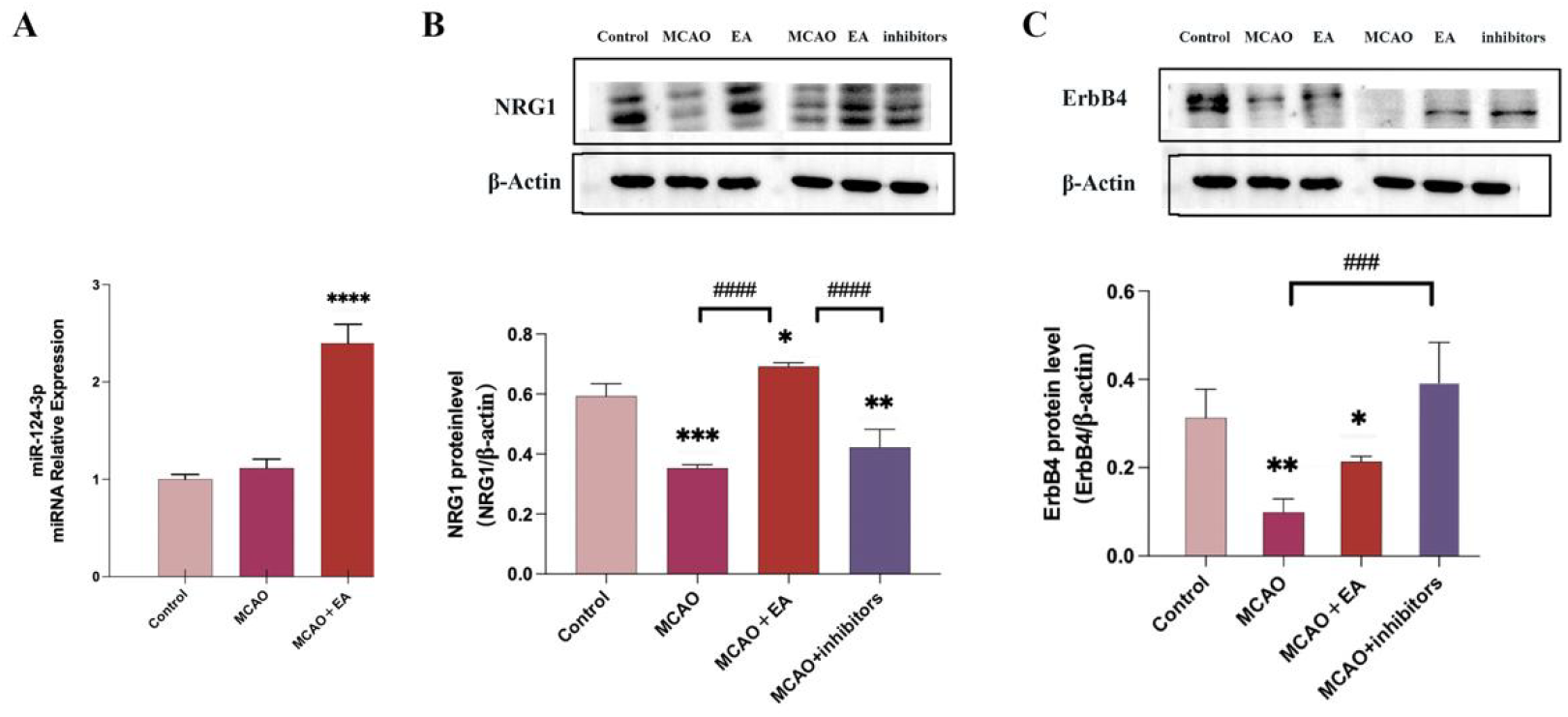

### 3.4 miR-124-3p alleviates MCAO/R-induced neuronal and axonal damage through the NRG1/ErbB4 signaling pathway

To investigate whether miR-124-3p alleviates neuronal and axonal damage by regulating GABA release through the NRG1/ErbB4 signalling pathway, we performed qPCR and Western blot analyses. qPCR results showed that compared with the model group, miR-124-3p levels were significantly upregulated in the electroacupuncture group (*P* < 0.05), whereas miR-124-3p expression was inhibited in the model plus miR-124-3p inhibitor group (Figure 4A). Western blot results showed that compared with the model group, protein expression of NRG1 and GABA was significantly upregulated in the electroacupuncture group (*P* < 0.05), while NRG1 protein expression was decreased in the model plus inhibitor group (*P* < 0.05) (Figure 4B). GABA expression showed an increasing trend in the inhibitor group (Figure 4C). These results suggest that miR-124-3p may alleviate neuronal and axonal damage by regulating GABA release through the NRG1/ErbB4 signaling pathway.

**Fig 4.**
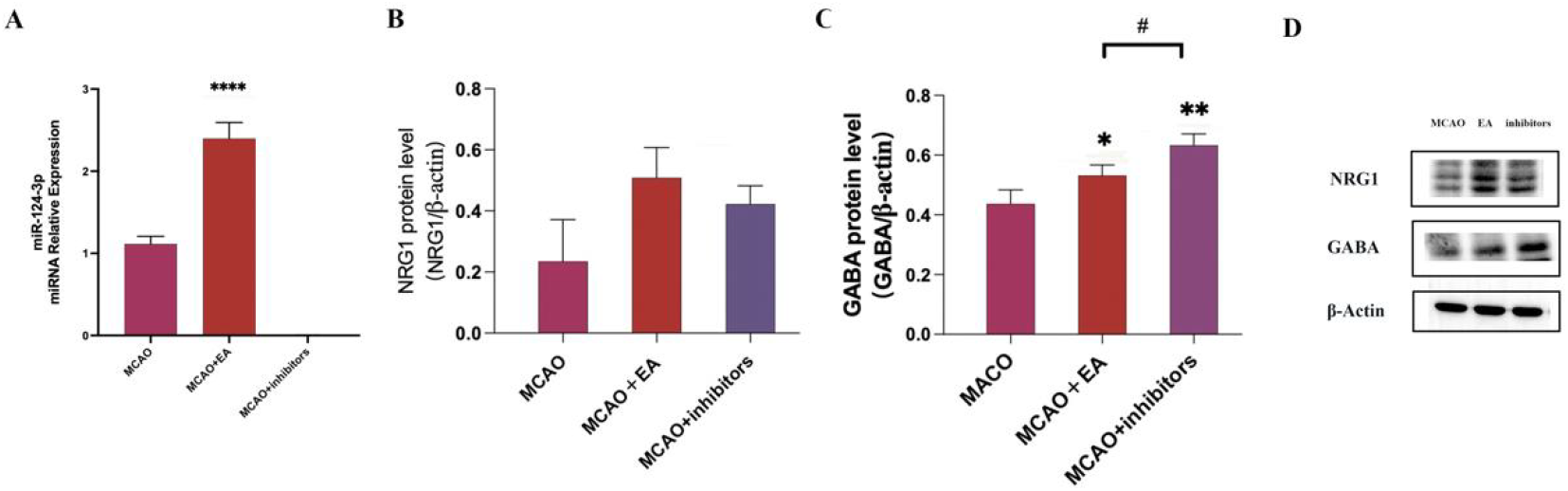

### 3.5 miR-124-3p directly targets and binds to the 3’UTR region of NRG1

The dual-luciferase reporter assay showed that the luciferase activity of the reporter gene containing the wild-type 3’UTR of NRG1 (NRG1 WT) was significantly decreased (P < 0.05) in the miR-124-3p mimic transfection group, whereas no significant change was observed in the luciferase activity of the reporter gene containing the mutant 3’UTR of NRG1 (NRG1 mut). These results confirmed that miR-124-3p specifically binds to the 3’UTR region of NRG1 mRNA, thereby regulating NRG1 expression at the translational level (Figure 5A and 5B).

**Fig 5.**
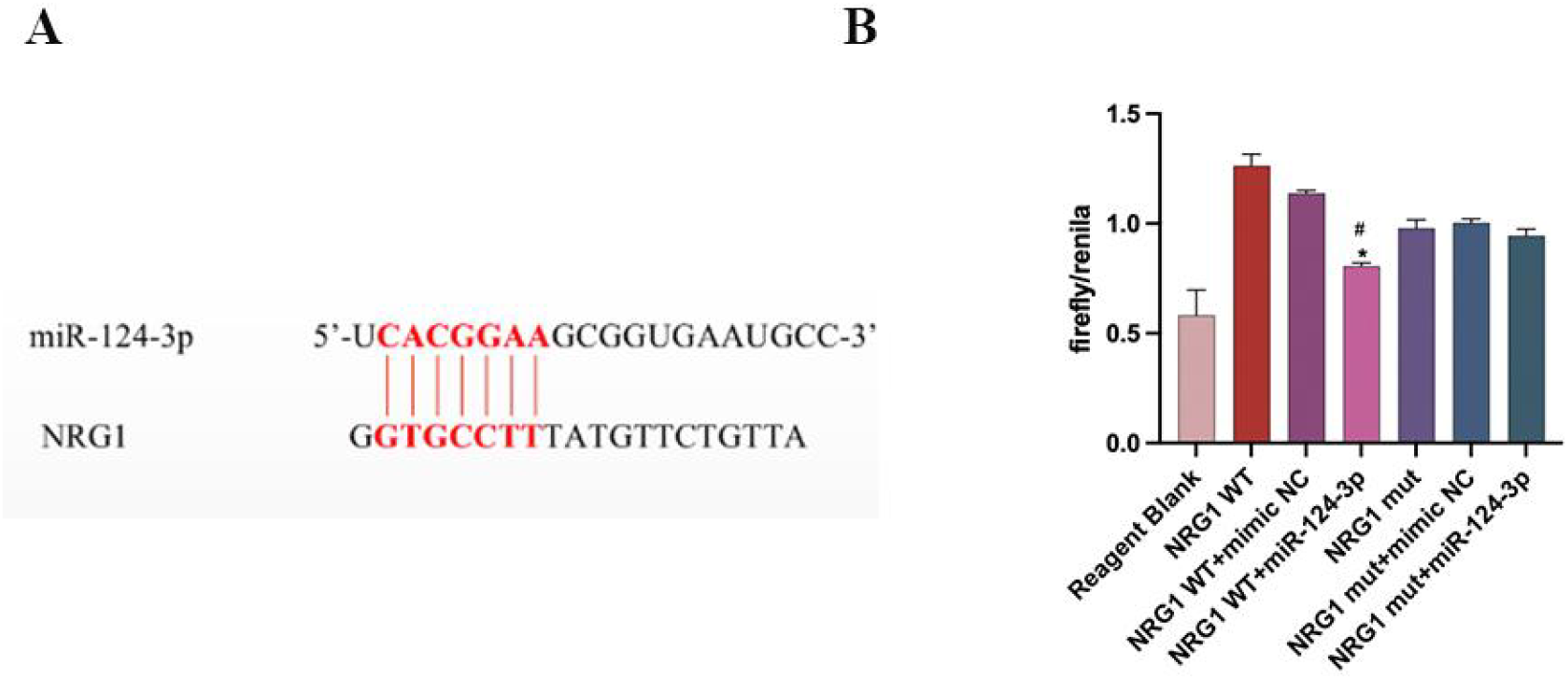

## 4 Discussion

In this study, we found that acupuncture improved motor function in MCAO/R model rats. The therapeutic effects may be associated with multi-neuroplasticity, as evidenced by acupuncture-induced increased expression of miR-124-3p, NRG1, ErbB4, and GABA in the hippocampus. Furthermore, miR-124-3p antagonist significantly eliminated the acupuncture-induced therapeutic effects, supporting this argument.

Stroke is a common cerebrovascular disease that can lead to numerous complications, with motor dysfunction being one of the primary symptoms^2,8,21^. Acupuncture, as a traditional therapeutic approach, has been shown to be effective in alleviating various central nervous system injuries^19,22^. ST36, located on the posterolateral aspect of the knee joint, is the most commonly used acupoint to improve symptoms in patients with motor dysfunction. Low-intensity acupuncture stimulation at ST36 facilitates nerve impulse conduction, induces neurotransmitter release and regulates brain function^23^. LI11, located at the lateral end of the elbow crease in the depression on the medial side of the radial wrist extensor, is the He-Sea point of the Large Intestine meridian of Hand-Yangming. The combined use of these two acupoints harmonizes meridian qi and blood circulation while relaxing tendons and collaterals, making them commonly used and effective acupoints for stroke treatment. This combination can better improve the symptoms of neurological deficits caused by ischemic stroke^24-26^.

Our animal behavioral studies confirmed the therapeutic effects of the electroacupuncture intervention. Zea Longa scores and pain threshold tests showed that rats in the electroacupuncture intervention group showed significant improvement in neurological deficits compared to the model group. Histopathological examination supported this finding. Using HE staining and Nissl staining of hippocampal neurons in MCAO/R rats, we observed that neuronal damage was significantly reduced after electroacupuncture treatment, with fewer abnormally shaped deep blue Nissl bodies. These results suggest that electroacupuncture treatment can mitigate MCAO/R-induced neuronal damage and ameliorate neuronal morphological abnormalities.

MicroRNAs play a crucial role in post-stroke recovery by regulating gene expression^27,28^. Previous studies have shown that miR-124-3p expression decreases significantly after ischemic brain injury, and restoring its expression can effectively promote neurological functional recovery^29,30^. Our results confirm the key role of miR-124-3p in electroacupuncture-promoted neural repair. Experimental findings demonstrated that electroacupuncture significantly upregulated miR-124-3p expression, while the use of a miR-124-3p antagonist inhibited the therapeutic effects of electroacupuncture. This finding is consistent with previous research on the important role of miR-124-3p in neural development and plasticity^31^.

We propose several potential mechanisms by which electroacupuncture upregulates miR-124-3p expression. First, electroacupuncture may activate neurotrophic factors such as brain-derived neurotrophic factor (brain-derived neurotrophic factor,BDNF) and nerve growth factor (nerve growth factor,NGF)^32,33^, which have been shown to regulate miR-124-3p transcription^31^. Second, electroacupuncture may facilitate calcium influx and activate calcium-dependent signaling pathways that influence miR-124-3p transcription factors^34^. In addition, electroacupuncture may indirectly enhance miR-124-3p expression by inhibiting pro-inflammatory cytokines such as TNF-α and IL-1β^35^, which are known to suppress miR-124-3p expression^30,36^. These proposed upstream mechanisms provide direction for future research, although further experimental validation is required.

Further investigation revealed that electroacupuncture significantly increased the expression levels of NRG1 and ErbB4. Dual-luciferase assays confirmed that miR-124-3p specifically binds to the NRG1 gene sequence, suggesting that miR-124-3p may exert its effects by regulating the NRG1-ErbB4 signaling pathway. After NRG1 binds to ErbB4, downstream signaling pathways can be activated, which play a crucial role in neuronal survival, axonal growth and synaptic plasticity^15^. We also found that electroacupuncture treatment significantly increased GABA expression^37^. As the primary inhibitory neurotransmitter in the central nervous system, GABA plays an essential role in maintaining neural circuit homeostasis. Our results suggest that this upregulation may be mediated by the NRG1-ErbB4 signaling pathway. These findings provide new evidence that electroacupuncture promotes recovery of motor function after stroke through the miR-124-3p/NRG1-ErbB4 pathway.

Extensive research shows that the brain rapidly recovers and reorganises its structure and function after stroke, with synaptic plasticity playing a crucial role in this process^38,39^. Several studies have confirmed that acupuncture can significantly increase the expression of synaptic plasticity-related proteins such as postsynaptic density protein 95, synaptophysin and NMDA receptors^32,40^. Our research suggests that electroacupuncture can regulate the miR-124-3p/NRG1-ErbB4/GABA pathway, which may work synergistically with these synaptic plasticity mechanisms. In addition, the NRG1-ErbB4 pathway has been shown to play an important role in the structural remodelling and functional regulation of pre- and postsynaptic membranes, modulating presynaptic neurotransmitter release and postsynaptic receptor clustering.

Compared with previous studies, this study has the following innovations. First, we established a complete miR-124-3p/NRG1-ErbB4/GABA signaling pathway, revealing a molecular cascade underlying the therapeutic effects of electroacupuncture, whereas previous studies mainly focused on individual molecules or single signaling pathways. Second, we directly confirmed the specific binding between miR-124-3p and NRG1 using a dual-luciferase reporter assay, providing clear evidence for the molecular targets of electroacupuncture treatment. These novel findings not only deepen the understanding of the mechanisms of electroacupuncture in stroke therapy, but also provide potential molecular targets for the development of new therapeutic strategies.

Despite yielding several significant findings, this study has limitations that should be addressed in future research. First, studies with larger sample sizes are needed to provide more robust experimental evidence for EA treatment of ischemic stroke. Second, the absence of real-time dynamic monitoring technology and functional imaging methods prevented comprehensive observation of immediate changes in molecular markers during electroacupuncture stimulation and its regulatory effects on brain functional networks. Although our study has established initial evidence for the interaction between miR-124-3p, the NRG1-ErbB4 signaling pathway, and GABA factors, the precise mechanisms by which miR-124-3p regulates the NRG1-ErbB4 signaling pathway remain unclear. Furthermore, the upstream regulatory mechanisms governing miR-124-3p following electroacupuncture stimulation require further investigation. Finally, the intervention period in this experiment was limited to 5 days, which restricted our ability to evaluate the long-term sustainability of the therapeutic effects.

In conclusion, this study elucidates the molecular mechanism by which electroacupuncture improves post-stroke neurological and motor dysfunction through the upregulation of miR-124-3p levels, which promotes the activation of the NRG1-ErbB4 signaling pathway and subsequently increases GABA release. These findings not only provide new theoretical evidence for electroacupuncture treatment of ischemic stroke, but also offer potential targets for the development of novel therapeutic strategies. Future research should focus on addressing the aforementioned limitations and further deepening our understanding of the mechanisms involved.

## 5 Conclusions

This study demonstrates that electroacupuncture at “Quchi” (LI11) and “Zusanli” (ST36) acupoints can improve motor function in rats with brain injury induced by MCAO/R. Mechanistically, electroacupuncture enhances the expression of miR-124-3p, which promotes the activation of the NRG1-ErbB4 signaling pathway, thereby facilitating GABA release and ameliorating post-stroke neurological and motor dysfunction. These findings provide new theoretical evidence for electroacupuncture treatment of post-stroke motor dysfunction. However, further research is needed to explore the mechanisms underlying electroacupuncture therapy for post-stroke motor dysfunction.

## 7 Author Contributions

DBY: Conceptualization, Investigation, Validation, Visualization, Writing – original draft, Writing – review and editing. PC: Methodology, Validation, Writing – original draft. DBY: Conceptualization, Formal Analysis, Software, Writing – original draft, Writing – review and editing. XTC: Data curation, Project administration, Supervision, Writing – original draft. YYL: Resources, Writing – original draft. FL: Methodology, Writing – original draft. NC: Writing – original draft. PC: Supervision, Writing – original draft. DBY and BS: Resources, Visualization, Writing – review and editing. BS: Project administration, Writing – review and editing.

## 9 Data availability statement

The original contributions presented in this study are included in this article/supplementary material, further inquiries can be directed to the corresponding authors.

## 10 Generative AI statement

The authors declare that no Generative AI was used in the creation of this manuscript.

## 11 Publisher’s not

All claims expressed in this article are solely those of the authors and do not necessarily represent those of their affiliated organizations, or those of the publisher, the editors and the reviewers. Any product that may be evaluated in this article, or claim that may be made by its manufacturer, is not guaranteed or endorsed by the publisher.

